# Memory specificity is linked to repetition effects in event-related potentials across the lifespan

**DOI:** 10.1101/2020.09.14.295972

**Authors:** Verena R. Sommer, Luzie Mount, Sarah Weigelt, Markus Werkle-Bergner, Myriam C. Sander

**Affiliations:** Center for Lifespan Psychology, Max Planck Institute for Human Development, Berlin, Germany; Department for Vision, Visual Impairments & Blindness, Faculty of Rehabilitation Sciences, TU Dortmund University, Germany

**Keywords:** episodic memory, memory specificity, repetition suppression, lifespan development, event-related potentials (ERP), electroencephalography (EEG)

## Abstract

The specificity with which past experiences can be remembered varies across the lifespan, possibly due to differences in how precisely information is encoded. Memory formation can be investigated through repetition effects, the common finding that neural activity is altered when stimuli are repeated. However, whether differences in this indirect measure of memory formation relate to lifespan differences in memory specificity has not yet been established. In the present study, we examined repetition effects in event-related potentials and their relation to recognition. During incidental encoding, children (aged 7–9 years), young adults (18–30 years), and older adults (65–76 years) viewed repeated object images from different categories. During subsequent recognition, we distinguished memory for the specific items versus the general categories. We identified repetition suppression in all age groups, and repetition enhancement for adults. Furthermore, individual item recognition performance comprising lure discrimination was positively associated with the magnitude of the neural repetition effects, which did not differ between groups, indicating common neural mechanisms of memory formation. Our findings demonstrate that neural repetition effects reflect the formation of highly specific memory representations and highlight their significance as a neural indicator of individual differences in episodic memory encoding across the lifespan.

## 1. Introduction

Memories of past events vary in their degree of specificity from very unique and detailed to more general and gist-like. Variability in memory specificity partly reflects how precisely incoming information has been encoded (McClelland and Rumelhart, 1985; Robin and Moscovitch, 2017). Forming precise memory representations is critical in many situations as it enables remembering specific details and avoiding confusion with similar information. This ability changes across lifespan development (cf. Ofen and Shing, 2013). Understanding how memory representations are formed is therefore crucial in the study of memory, particularly with regard to differences in memory competences across the lifespan.

One way to study the formation of neural representations is to examine brain activity in response to repeated stimulus input (Helen C. Barron et al., 2016; Grill-Spector et al., 2006). If the neural responses differ between first and repeated encounters, which is called repetition effect, this indicates that an internal representation of the repeated stimulus has been formed (Rugg and Doyle, 1994). Indeed, such repetition effects are widely observed, and potentially they can have two different directions: Repetition *suppression* describes the finding that neural activity evoked by a repeated stimulus is *reduced* (Helen C. Barron et al., 2016; Grill-Spector et al., 2006). For example, Stefanics et al. (2018) identified three time windows (86– 140, 322–360, and 400–446 ms after stimulus onset) in which repeated stimuli compared to stimuli that were shown for the first time showed reduced amplitudes in event-related potentials (ERPs) over occipital, temporoparietal, and frontotemporal regions. In addition, *increased* activity (repetition *enhancement)* has also been reported, especially in the ERP literature (Desimone, 1996; Doniger et al., 2001; Nagy and Rugg, 1989; Rugg et al., 1995), but also in functional magnetic resonance imaging (fMRI; for a review, see Segaert et al., 2013). For instance, Stefanics et al. (2018) also found enhanced ERP amplitudes (320–340 ms) for repeated stimuli at occipitotemporal electrodes. Thus, repetition effects manifest as repetition suppression and / or repetition enhancement effects, both in concert or in isolation. However, independent of their direction, repetition effects are indicative of the formation of a neural representation of the stimulus. Importantly, repetition effects vary with regard to particular *properties* of neural representations such as their specificity, i.e., the precision with which a certain content (and not other, e.g., similar content) is represented by neural activity (cf. Barron et al., 2016b; Koolschijn et al., 2019; Schacter et al., 2004). For instance, Jiang et al. (2006) modulated the similarity of face images by morphing and showed that moderately similar faces elicited smaller fMRI repetition suppression effects than identical faces, which was related to face discrimination performance. These findings suggest that repetition effects can track the specificity of neural representations (Gilaie-Dotan and Malach, 2007; Grill-Spector et al., 2006; Lueschow et al., 1994).

However, until now, the repetition effect has been mainly implicated as the neural correlate of implicit memory (namely repetition priming: Doniger et al., 2001; Gotts et al., 2012; Henson, 2003; Wiggs and Martin, 1998), and only rarely been associated with explicit memory performance (cf. Nagy and Rugg, 1989). Those few studies yielded a mixed pattern of results: On the within-person level, i.e., in terms of ‘subsequent memory effects’ (Paller and Wagner, 2002), Turk-Browne et al. (2006) identified larger repetition effects for subsequently remembered scenes compared to forgotten scenes. This result would suggest that the size of the repetition effect is indicative of successful memory formation. However, other studies found no relation (Rugg, 1990; Ward et al., 2013) or even a negative relation between neural repetition effects and explicit memory performance, meaning that a larger effect was associated with poorer memory (Wagner et al., 2000; Xue et al., 2011). On the between-person level, individual differences in fMRI repetition suppression were shown to be related to individual differences in memory performance and thus linked to memory deficits in particular groups only, such as in participants with autism spectrum conditions (Ewbank et al., 2017) or mild cognitive impairments and Alzheimer’s disease (Pihlajamäki et al., 2011). Given that previous findings are mixed, the current study addresses the question whether repetition effects can be leveraged as a useful indicator for successful episodic memory formation within and across different populations, for example, age groups.

Memory abilities vary across the lifespan, especially the ability to remember specific details versus the general gist of an episode (Graf and Ohta, 2002; Hultsch et al., 1998). The balance of being able to discriminate among similar events (pattern separation) and the ability to abstract and generalize between events (pattern completion) is critical for successful and adaptive behavior (McClelland and Rumelhart, 1985; Wiltgen and Silva, 2007; Xu and Südhof, 2013). Across childhood, the capability to differentiate similar from old information increases (Ngo et al., 2018; Rollins and Cloude, 2018). According to these findings, children’s memories develop from more general to more specific which has been associated with hippocampal maturation (Keresztes et al., 2018, 2017). However, at the same time, children’s memories have also shown to rely more on specific perceptual properties than on abstract/semantic knowledge that is still developing, making them *less* susceptible to confuse conceptually similar information (Brainerd et al., 2002, 2008; Howe, 2008; Ofen and Shing, 2013; Sloutsky and Fisher, 2004; cf. fuzzy trace theory, Brainerd and Reyna, 1990). At the other end of the lifespan, older adults have shown to rely more on general concepts and similarities, leading to more false memories (Fandakova et al., 2018; Koutstaal and Schacter, 1997; Schacter et al., 1997), especially concerning the discrimination of similar information (Dennis et al., 2007; Stark et al., 2015, 2013; Toner et al., 2009).

Developmental differences in memory abilities are accompanied by age-dependent changes in the underlying brain structures and functions during maturation and senescence (Li et al., 2004; Ofen and Shing, 2013; Sander et al., 2020a; Shing et al., 2010; Van Petten, 2004). Specifically, functional differences during encoding may influence the specificity of mnemonic contents and may thus account for performance differences across the lifespan (Bowman et al., 2019; Fandakova et al., 2019; Morcom, 2015; Park et al., 2013; Sander et al., 2020b). In line with this, studies measuring neural specificity as the discriminability of BOLD activation patterns have shown a positive relation between the specificity of these patterns during encoding and memory performance in young and older adults (Kobelt et al., 2020; Koen et al., 2019) as well as in children (aged 7–12 years, (Fandakova et al., 2019). Similarly, studies using neural repetition effects to indicate how precisely information is represented (see above; Jiang et al., 2006) found that the specificity with which faces were represented in face-selective brain regions changes across the lifespan (cf. Nordt et al., 2016): with lower specificity in children (5–12 years) than young adults (Natu et al., 2016), and in older adults (61–72 years) than in young adults (Goh et al., 2010). In both studies, neural repetition effects were associated with differences in face discrimination performance. Thus, piecing together evidence from different lines of research addressing the two ends of the lifespan separately, the available evidence points to an improvement of how precisely information is encoded from childhood to adulthood followed by a decline during aging. However, age-comparative studies including both children and older adults that directly test the association between repetition effects and memory abilities, particularly memory specificity, are lacking.

In the current EEG study, we therefore investigated whether neural repetition effects were directly associated with the specificity of later recognition by studying ERP repetition effects in response to repeated object images in children (7–9 years), young adults (18–30 years), and older adults (65–76 years). We investigated how individual differences in repetition effects (i.e., the magnitude of suppression and/or enhancement effects) were related to the ability to recognize the presented objects. We hypothesized that potential lifespan age differences in memory performance might be paralleled by age group differences on the neural level. During recognition, exact repetitions, similar lures, and entirely new items were presented (cf. Stark et al., 2019), allowing for a dissociation of mere category memory from precise exemplar memory. This enabled us to investigate whether repetition effects reflected general concept or detailed item-specific encoding.

## 2. Materials and Methods

### 2.1 Participants

A total of 23 healthy children, 41 healthy young adults, and 61 healthy older adults participated in the EEG study. Further collection of children’s data had to be aborted due to the COVID-19 pandemic. We excluded one child and three older adults due to low performance in the vigilance task during encoding (see Behavioral performance) and one child, two young adults, and two older adults due to missing or noisy EEG data. One additional older adult was excluded based on low performance in the dementia screening (see below). The remaining 21 children, 39 young adults, and 55 older adults are included in the analyses presented here (see demographic information in Table 1).

**Table 1.** Demographics

| Group | Collected<br>*N* | Included<br>*N* | Female<br>*N* | Male<br>*N* | Age *M*(*SD*) | Age<br>range | Education<sup>1</sup><br>*M*(*SD*) |
| --- | --- | --- | --- | --- | --- | --- | --- |
| Children | 23 | 21 | 15 | 6 | 8.38(0.74) | 7–9 | N/A |
| Young adults<br>(main) | 41 | 39 | 23 | 16 | 24.77(3.30) | 18–30 | 15.04(4.46) |
| Young adults<br>(control) | 23 | 23 | 13 | 10 | 23.38(2.65) | 19–29 | N/A |
| Older adults | 61 | 55 | 27 | 28 | 70.15(3.39) | 65–76 | 13.24(3.24) |

*M* = mean, *SD* = standard deviation, <sup>1</sup> Years incl. higher education

School children (7–9-year-olds) were selected for the children’s age group such that they would have already experience with longer phases of concentrating on a given task (approximately 45 minutes). Since in this age range recognition memory (Golarai et al., 2007; Rollins and Cloude, 2018) and perhaps even memory specificity (Ngo et al., 2018) may already be very advanced, we did not include children older than 9 years.

Children were recruited from a participant database at the Developmental Neuropsychology Lab at the Ruhr-Universität Bochum (RUB), Germany. Adults were recruited from a participant database at the Max Planck Institute for Human Development in Berlin, Germany, and through advertisements.

All participants were native German speakers, had normal or corrected-to-normal vision, no history of psychiatric or neurological disease, and no use of psychiatric medication. We screened older adults with the Mini-Mental State Examination (MMSE; Folstein et al., 1975). One had a score below the threshold of 24 points (namely 21) and was excluded. The remaining older adults reached a mean score of 28.95 (SD = 1.21, 25–30 points).

To control for the different task versions (see below), behavioral performance of 23 additional young adults on the shorter children’s task was assessed at RUB.

All experiments were approved by the respective local ethics committee. All participants and the children’s parents gave informed consent to taking part in the experiment and all participants and/or their parents could abort the study at any time without giving reasons.

### 2.2 Experimental design

Data for this study were collected at two different sites. Children were tested at the Developmental Neuropsychology Lab at the Ruhr-Universität Bochum, Germany, and adults were tested at the Center for Lifespan Psychology at the Max Planck Institute for Human Development in Berlin, Germany. Some experimental procedures, such as collected covariates, also differed for children and adults (see Table S1). However, the same visual object encoding and recognition tasks were at the core of the study (see Figure 1), with children performing a shorter version of the tasks, namely only the first half. Other than that, the selection and order of stimuli in both tasks were identical for all participants.

**Figure 1.**
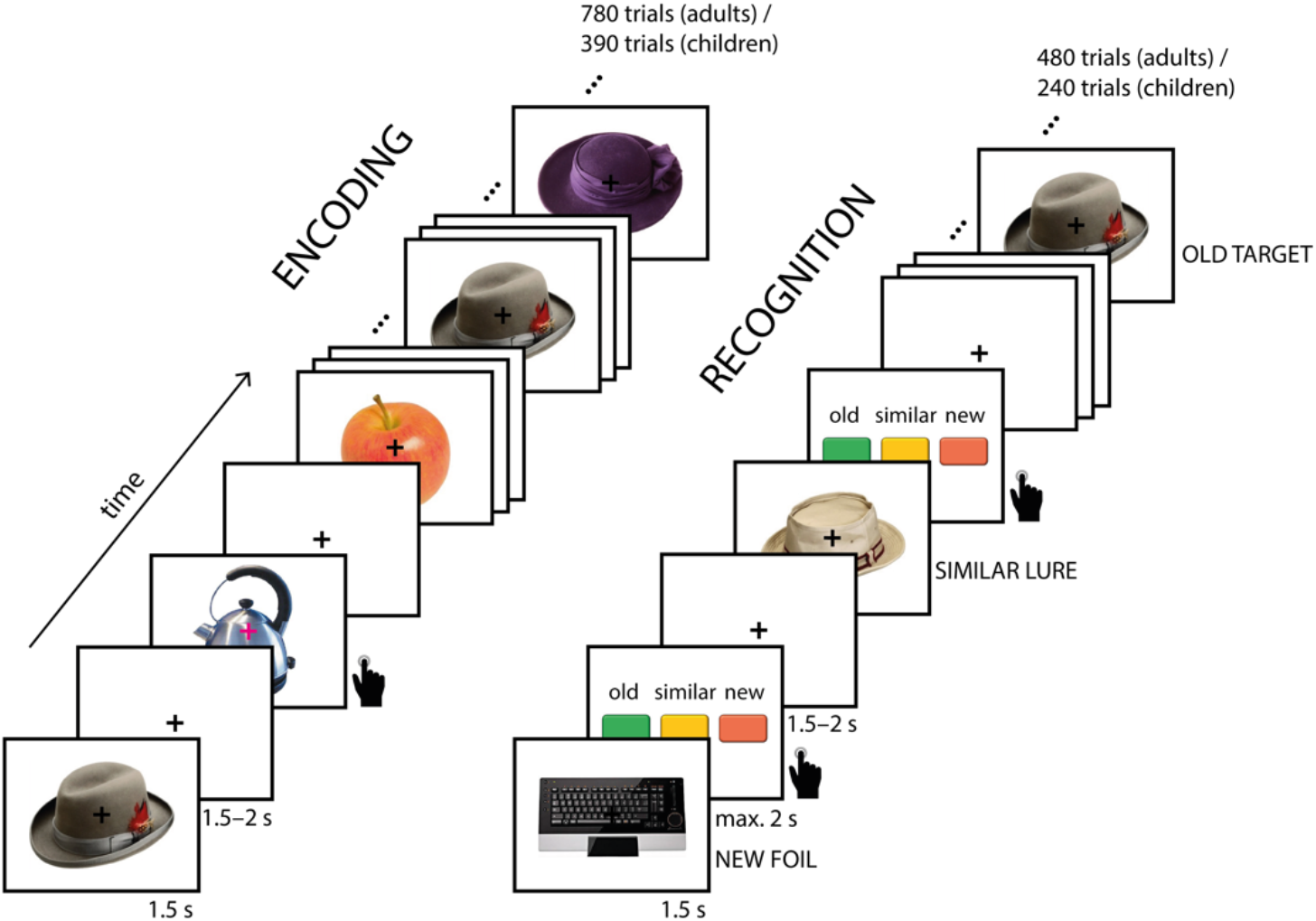
Overview of the experimental paradigm. In the encoding task, objects were sequentially presented, and subjects were asked to press a button whenever the fixation cross changed color. During recognition, objects were sequentially presented, and subjects were asked to decide for each object whether it was old, similar, or new. Each item was shown only once.

During encoding and recognition, participants viewed color photographs of objects from different categories (e.g., hats, trees, guitars). The pictures were partly selected from a published stimulus set (“Massive Memory” Object Categories; https://konklab.fas.harvard.edu; Konkle et al., 2010) and partly obtained from the internet. The stimuli were scaled to 500 × 500 pixels and all text was removed using Adobe Photoshop CC 2017. Stimuli were presented using Psychtoolbox (Psychophysics Toolbox, RRID: SCR_002881) for MATLAB (The MathWorks Inc., Natick, MA, USA; RRID: SCR_001622). During both tasks, EEG was acquired.

#### 2.2.1 Encoding task

In the following, we describe the task paradigm for the adults, followed by the specific differences for the children’s sample. During the encoding task (see Figure 1), adult participants viewed objects from 80 different categories. Participants were not instructed to memorize the stimuli and did not know that their memory would be tested later on. Stimuli were presented successively on a white background on the center of the screen. A central fixation cross was superimposed on the objects and remained on the screen throughout the task. Stimulus duration was 1500 ms with an inter-stimulus-interval jittered between 1500 and 2000 ms. Subjects were instructed to attend to the objects but to fixate on the cross in order to minimize eye movements. To ensure that participants attended to each trial, they performed a target detection task in which they were asked to press a button whenever the fixation cross changed its color from black to magenta. The target trials appeared in approximately 7% of all trials, that is 54 times in total, with 1–25 (on average 14.7) trials between two target trials. The color change occurred 100–500 ms after onset of the stimulus. The target trials showed objects from additional categories and were excluded from further analyses.

After the first half of the trials, adult participants could take a voluntary break. To maintain motivation for the following trials, participants received visual feedback on how many color changes they had successfully detected so far. The two halves of the encoding task were independent of each other in that categories from the first half were not part of the second half.

The 80 object categories were equally divided into four conditions that differed with respect to (a) the number of different presented exemplars from one category (either two or four), and (b) the number of exemplar repetitions (either two or four times). Details on the four conditions can be found in the supplement. For the purpose of the current analysis, we collapsed across all conditions and focused on the comparison of the first and second presentation of an object, independent of its condition.

The stimulus order was pseudorandomized with the restriction that 3–10 stimuli (from other categories) appeared between repetitions of the same item, with the same mean distance for all conditions. Furthermore, at least five other items were presented after the last repetition of a category exemplar before the next exemplar of that category was shown. In this way, exemplars from the same category were not presented interleaved. A total of 720 experimental trials was presented. In addition to these experimental trials, 54 target trials (see above) and 6 filler trials were presented. Filler trials were needed when the distance restrictions of the pseudorandomization could not be met otherwise. Like target trials, filler trials showed objects from additional categories and they were excluded from further EEG analyses. Since only first and second object presentations are included in the current analyses, only 320 of the 720 experimental trials were analyzed here (160 first and 160 second presentations).

A shorter version of the task was used for the children. They performed only the first half of the adult task, comprising 40 distinct object categories presented in a total of 360 experimental trials, plus 27 target trials, and 3 filler trials. From these total number of trials, 160 trials were analyzed here (80 first and 80 second object presentations). In contrast to the adults, children had more breaks, namely every 85 trials (approximately every 4 minutes). The breaks were voluntary and self-paced.

#### 2.2.2 Recognition task

After a 15-minute break, a surprise recognition task followed (see Figure 1). Again, object images were presented successively on a white screen and with a black fixation cross on top. The presented objects were either “old,” i.e., identical to an item of the encoding task (targets), “similar,” i.e., of a category that was already presented in the encoding task but a different exemplar (lures), or “new,” i.e., of a novel category (foils). Stimulus duration was again 1500 ms. After each stimulus presentation, participants were asked to indicate whether the object was “old,” “similar,” or “new” by pressing one of three buttons on a button box. The response was self-paced but limited to 2000 ms. Thereupon, the fixation cross was shown for 1500– 2000 ms before the next stimulus item appeared.

For adults, the task consisted of 160 targets, 160 lures, and 160 foils, resulting in a total of 480 trials and a chance level of 33%. The targets contained the same 80 object categories that were presented in the encoding task, but for each category, only two exemplars were presented in the recognition task. Two more novel exemplars from the same categories were presented as similar lures. The foils consisted of two exemplars from each of the 80 novel object categories. The stimuli were pseudorandomized with the restriction that there were at least three trials between items from the same object category, and maximally four stimuli with the same correct response (“old,” “similar,” “new”) were presented in a row. Targets that were presented in the first/second half of the encoding task were also tested in the first/second half of the recognition task, respectively.

The children only performed the first half of the recognition task, i.e., the first 240 trials, containing 80 targets, 80 lures, and 80 foils. In contrast to the adults who had no breaks in the recognition task, the children could again take a self-paced break every 85 trials.

### 2.3 Analysis of behavioral data

#### 2.3.1 Encoding task

Performance in the encoding task was measured as the proportion of correctly detected color changes of the fixation cross.

#### 2.3.2 Recognition task

In the recognition task, participants were asked to indicate for each presented object whether it was “old” (target), “similar” (lure), or “new” (foil). One way of quantifying memory performance is to calculate the proportion of correct answers: *Pr(“old” | target) + Pr(“similar” | lure) + Pr(“new” | foil).* However, the two respective incorrect responses for each item class can be evaluated as not equally incorrect but as reflecting the level of memory specificity. For example, incorrectly responding “similar” to a target is less precise than responding “old,” but it shows that the general category has been correctly identified as known, whereas a “new” response would reflect no memory for either the item or category.

On this basis, we computed performance measures that focus on different levels of specificity, namely on precise item memory and more general category memory (see Figure 2). These measures are equivalent to the classic performance measure *d’* in old/new recognition tasks that assess correctly recognized old items (hits) corrected for falsely recognized new items or the tendency to respond “old” (false alarms). The current recognition task comprised three response options with graded precision. Thus, different levels of memory specificity can be measured by scoring different responses as correct versus incorrect.

**Figure 2.**
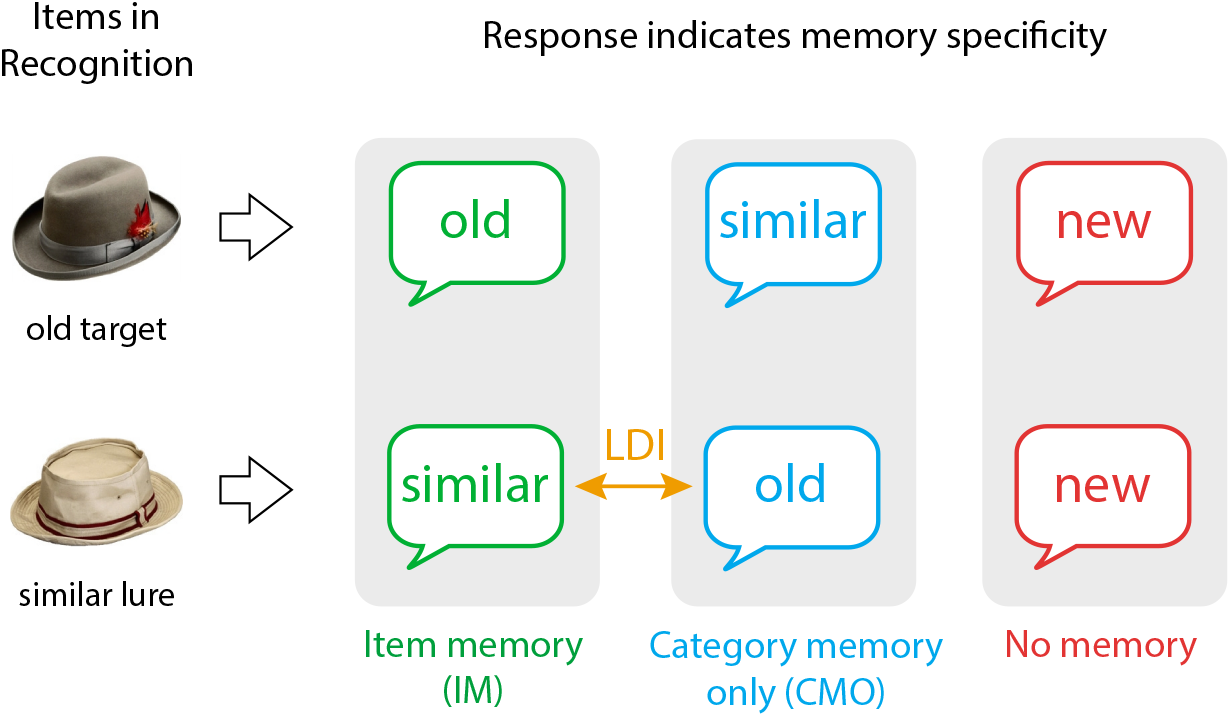
Illustration of memory specificity measures. Correctly identifying targets as old and lures as similar reflects specific item memory (IM; green), incorrectly identifying targets as similar and lures as old reflects mere category memory (CMO; blue), and incorrectly identifying targets and lures as new reflects no memory (red). The difference between identifying lures as similar and mistaking them as old is the lure discrimination index (LDI; yellow). The measures of IM and CMO are furthermore corrected for the tendency to respond “old” or “similar” (not depicted).

##### Item memory (IM)

Only if specific details of the objects were encoded, were participants able to distinguish previously presented targets from similar lures and vice versa. In this measure, only responses of correct target memory and lure detection were scored as accurate, corrected for both unspecific category memory (see below) as well as the general tendency to respond “old” or “similar.”

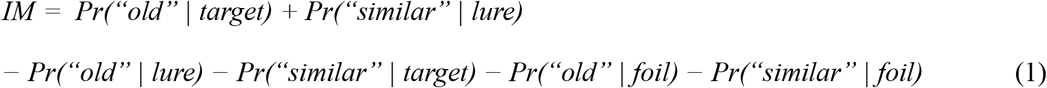

In terms of probability, since IM is corrected for four incorrect response combinations while only two combinations are counted as correct, it is more likely to give an incorrect than a correct response. Because of this imbalance of correct and incorrect response probabilities, the chance level of IM is below 0, namely –0.22. To have all measures on the same scale, we adjusted IM (by adding 0.22) such that it denotes the deviation from its chance level.

##### Category memory only (CMO)

This performance measure captures the ability to successfully remember the general category of presented objects but a failure to distinguish individual exemplars, corrected for the general response bias. Thus, CMO is a measure of *mere* category memory, and not category memory in general, which would include item memory as well.

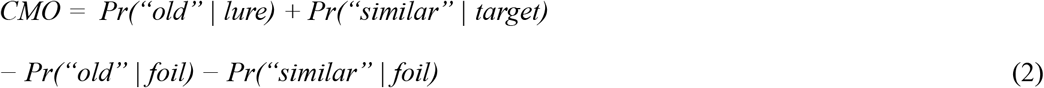

These performance measures emphasize different degrees of memory specificity. For CMO, participants show only a sense of familiarity for the object categories, whereas high IM performance requires remembering specific object details in order to recognize the exact exemplar and to reject a similar lure.

##### Lure discrimination index (LDI)

One component of item memory is lure discrimination. The ability to discriminate lures has been assessed separately in a number of studies and associated with hippocampal pattern separation/completion and age differences therein (e.g., Keresztes et al., 2017; Morcom, 2015; Ngo et al., 2018; Stark et al., 2013; Toner et al., 2009; Yassa et al., 2011). The measure reported here computes the difference between correct lure detection and the susceptibility to mistake lures as old (cf. Ngo et al., 2018; Rollins and Cloude, 2018; Toner et al., 2009).

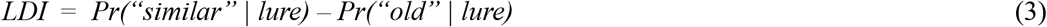

### 2.4 EEG recording and preprocessing

EEG was recorded during the object encoding and recognition tasks. In adults, EEG was recorded continuously with BrainVision amplifiers (BrainVision Products GmbH, Gilching, Germany) from 61 Ag/AgCl electrodes embedded in an elastic cap (EASYCAP GmbH, Herrsching, Germany). Two additional electrodes were placed at the outer canthi and one below the left eye to monitor eye movements (horizontal and vertical EOG). During recording, all electrodes were referenced to the right mastoid electrode, and the left mastoid electrode was recorded as an additional channel. Electrode AFz served as ground. The EEG was recorded with a pass-band of 0.1 to 250 Hz and digitized with a sampling rate of 1000 Hz. During preparation, electrode impedances were kept below 5 kΩ.

In children, EEG was recorded with BrainVision amplifiers from 64 active Ag/AgCl electrodes in an ActiCAP cap with electrode FCz as reference. Everything else was identical to the adults.

For all age groups, the same EEG data preprocessing pipeline was used, except that adults’ EEG and eye-tracking data were first integrated using the Eye-EEG toolbox (Dimigen et al., 2011; RRID: SCR 012903). EEG preprocessing was performed with the EEGLab (Delorme and Makeig, 2004; RRID: SCR 007292) and Fieldtrip (Oostenveld et al., 2011; RRID: SCR 004849) toolboxes as well as custom MATLAB code (The MathWorks Inc., Natick, MA, USA; RRID: SCR 001622). For analyses, EEG data were demeaned, re-referenced to an average reference, downsampled to 250 Hz, and band-pass filtered (0.2–125 Hz; fourth order Butterworth). Data were visually screened for excessive muscle artifacts, followed by an independent component analysis (ICA) to identify components related to eye, muscle, and cardiac artifacts (Jung et al., 2000). Microsaccadic artifacts were detected by using an algorithm for the correction of saccade-related transient spike potentials (COSTRAP; Hassler et al., 2011). Artifact components were automatically detected, visually checked, and removed from the data. Following the FASTER procedure (Nolan et al., 2010), automatic artifact correction was then performed for any residual artifacts. Channels excluded prior to ICA were interpolated after artifact correction with spherical splines (Perrin et al., 1989). The mean number of interpolated channels was 2.1 (maximum = 5). Finally, 10% of the remaining trials were again visually screened to determine successful cleansing. For further analysis, data epochs of 5.5 s were extracted from –2 s to 3.5 s with respect to the onset of the object presentation. After preprocessing, the mean total number of encoding trials was 328.62 for children (range = 291–352), 680.21 for young adults (range = 590–705), and 686.75 for older adults (range = 591–711).

### 2.5 EEG analysis

The current paper only covers analysis of the EEG recorded during encoding. Furthermore, since there were no interactions between age and item manipulations during encoding, we largely disregard the encoding conditions and do not separate the trials accordingly in the following analyses (see Supplements). We computed ERPs by averaging the time-locked EEG data over the respective trials, separately for each electrode (see Supplementary Figure S1 for ERPs at selected electrodes). For ERP group comparison, we applied an absolute pre-stimulus baseline correction (200 ms before onset). To examine repetition effects, we calculated ERPs separately for the first and second presentation of the object items.

### 2.6 Statistical analysis

#### 2.6.1 Memory performance

For each age group, we tested whether participants performed better than chance in the recognition task by contrasting the proportion of correct responses to chance level (0.33) (one-sample *t*-tests). We tested for age differences in all reported behavioral measures by conducting one-way ANOVAs contrasting children, young adults, and older adults, followed by post-hoc *t*-tests.

#### 2.6.2 Event-related potentials

On the first (within-subject) level, we compared the ERPs of every encoding trial in which an object item was presented for the first time with the ERPs of the respective second presentation using two-sided paired sample *t*-tests.

On the second (between-subject) level, the individual *t*-maps of the first-level analysis were contrasted against zero using two-sided independent samples *t*-tests. We controlled for multiple comparison by conducting non-parametric, cluster-based, random permutation tests (Fieldtrip toolbox; RRID: SCR 004849; Maris and Oostenveld, 2007; Oostenveld et al., 2011; see also Fields and Kuperberg, 2019). All scalp electrodes (60 for adults, 64 for children) and the trial time points from −0.1 to 1.6 s relative to stimulus onset were included in the analysis. Clusters were formed by grouping neighboring channel × time samples with a *p*-value below 0.05 (spatially and temporally). The respective test statistic was then determined as the sum of all *t*-values within a cluster. The Monte Carlo method was used to compute the reference distribution for the summed cluster-level *t*-values. Samples were repeatedly (10,000 times) assigned into two classes and the differences between these random classes were contrasted with the differences between the actual two conditions (e.g., first and second presentation). For every permutation the *t*-statistic was computed, and the *t*-values summed for each cluster. This was done separately for each age group to reveal the group-specific repetition effects.

On the third (between-group) level, we tested whether the identified effects differed between age groups. For this, we extracted and averaged the individual *t*-values from the first-level analysis within the time–electrode clusters identified in the second-level analysis. For all group comparisons, *t*-values were transformed to *z*-values. For each repetition effect, these values were entered into a one-way ANOVA contrasting children, young adults, and older adults, or underwent a two-sample *t*-test if only two groups were compared.

We tested whether the identified clusters may be different reflections of the same repetition suppression/enhancement effect by correlating the individual effect sizes, i.e., the extracted mean *t*-values, across subjects (Pearson correlation). If clusters correlated highly (*r* > 0.7), we pooled them together as one composite effect by averaging across the individual mean *t*-values.

#### 2.6.3 Brain–behavior relationship

We tested for a relationship between ERP repetition effects (first versus second presentation) and memory performance by modelling their correlations using maximum likelihood estimations (in Ωnyx 1.0-1010; von Oertzen et al., 2015). Specifically, the correlations of the individual mean *z*-transformed *t*-values extracted from the identified time–electrode clusters (see above) and the individual measures of memory performance were modelled (a) separately for each age group, and (b) across all age groups. Since we observed overall age differences in both effect sizes and memory performance, we standardized both measures within groups in order to test for the brain–behavior association independent of overall age differences. We tested whether the models (a and b) showed significantly different model fits using likelihood-ratio (chi-squared) tests. Significant differences between the model fits (H1) would show that the model allowing for group-specific correlations (a) explains the data better, indicating that the correlations differed between groups, which would then require follow-up analyses of the pairwise differences. In turn, if the model fits did not differ significantly (H0), this would indicate that the brain–behavior relationship was not different between groups.

### 2.7 Code and data accessibility

Code and summarized data will be made available upon publication.

## 3. Results

### 3.1 Behavioral performance

#### 3.1.1 Encoding task

During encoding, the participants’ task was to detect when the fixation cross changed its color. We expected close-to-ceiling performance when participants attended to the stimuli and therefore excluded everyone who detected less than 80% of the respective trials. One child (0.67 hit rate) and two older adults (0.72 and 0.50 hit rate, respectively) were excluded based on this criterion. The remaining participants had a mean (±SD) hit rate of 0.98 (0.03) (children), 0.99 (0.02) (young adults), and 0.96 (0.04) (older adults).

#### 3.1.2 Recognition task

In the recognition task, participants were asked to decide for each presented object whether it was “old” (target), “similar” (lure), or “new” (foil). The mean (±SD) proportion of correct responses was 0.63 (0.17) for children, 0.57 (0.15) for young adults, and 0.56 (0.15) for older adults (chance level: 0.33). On average, subjects of all age groups performed better than chance (children: *t*(20) = 17.47, *p* < 0.001; young adults: *t*(38) = 18.09, *p* < 0.001; older adults: *t*(54) = 19.13, *p* < 0.001; one-sample *t*-tests of proportion correct responses against chance level). The mean number of “old”, “similar”, and “new” responses to targets, lures, and foils is shown in Figure 3.

**Figure 3.**
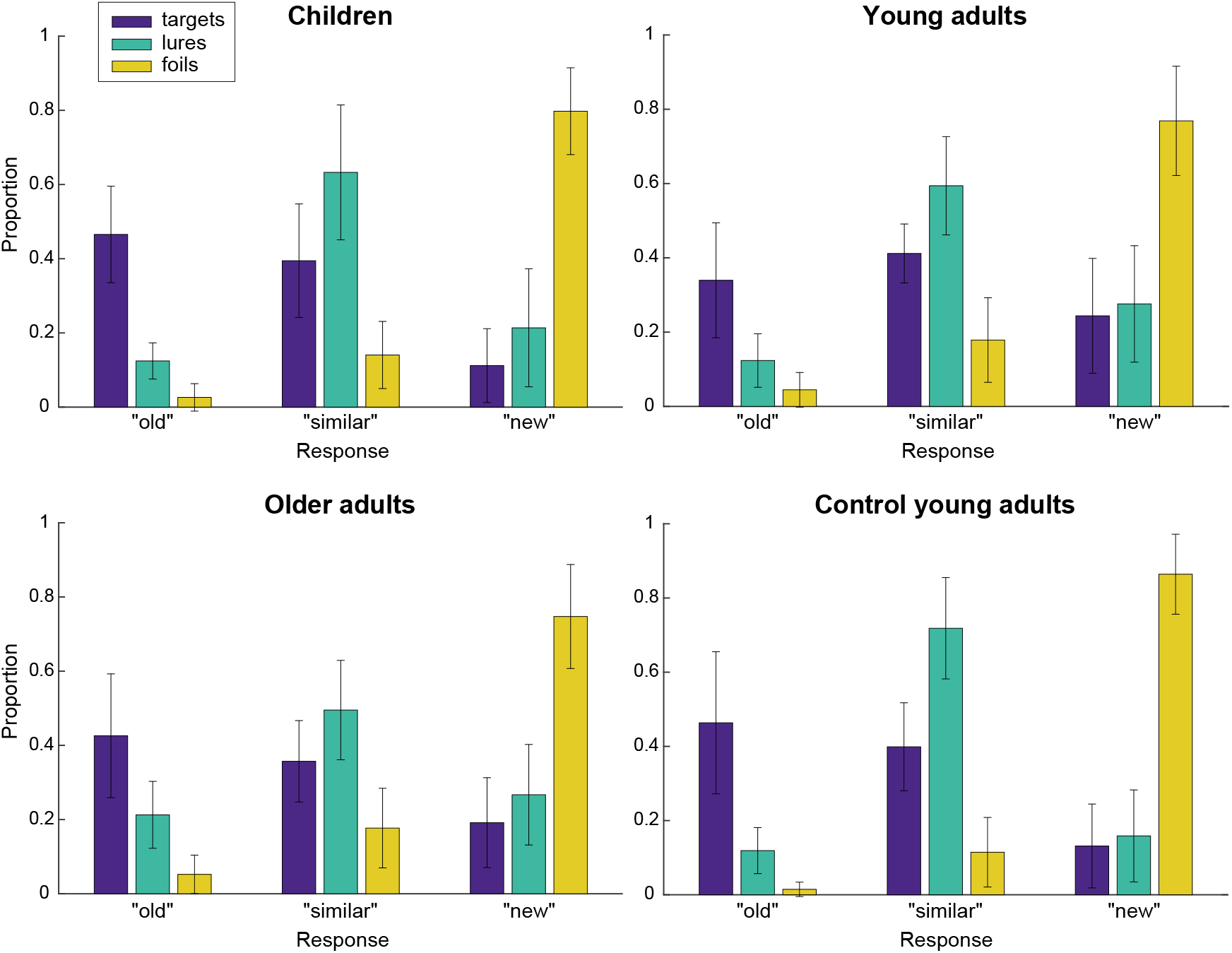
Mean proportion of responses (“old”, “similar”, “new”) to targets, lures, and foils (color-coded) for 21 children, 39 young adults (EEG group), 55 older adults, and 23 additional control young adults (only behavioral). Children and control young adults performed the shorter task version. Error bars denote standard deviation of the mean.

Age differences in memory specificity. Highly specific item memory (IM; Eq. (1); Figure 4A) showed significant age differences (*F*(2,112) = 7.76, *p* < 0.001; one-way ANOVA). Post-hoc *t*-tests revealed higher performance of children compared with young adults (*t*(58) = 3.27, *p* = 0.002) as well as older adults (*t*(74) = 3.81,*p* < 0.001), but no differences between the adult groups (*t*(92) = 0.78,*p* = 0.435). In contrast, correct category in combination with incorrect item memory (CMO; Eq. (2); Figure 4B), showed no age differences (*F*(2,112) = 1.21, *p* = 0.301). That is, the age groups differed in their specific exemplar memory but not mere category memory.

**Figure 4.**
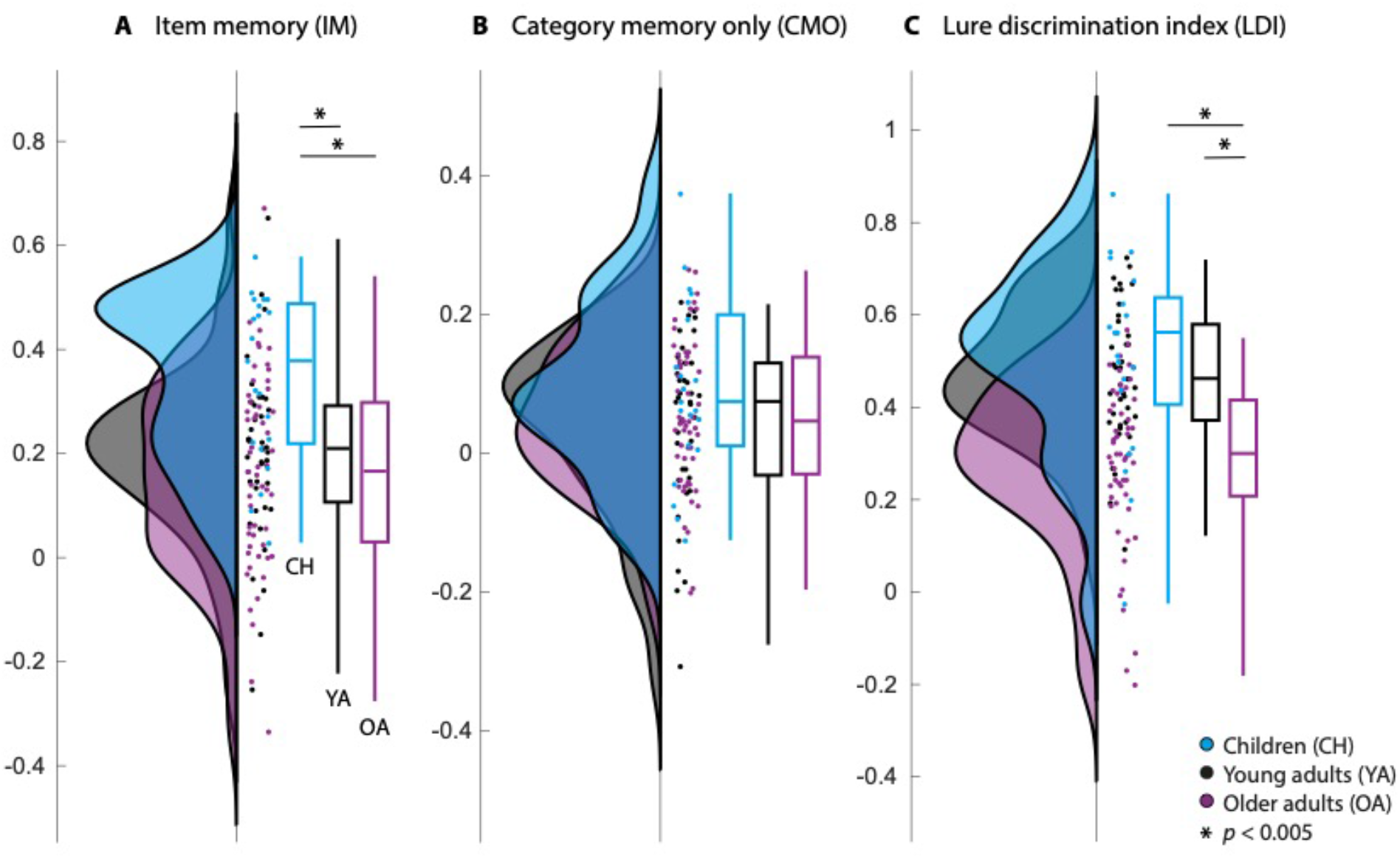
Behavioral performance of the children (CH; blue), young adults (YA; black), and older adults (OA; purple). **A.** Correct item memory (IM). **B.** Category memory only (CMO). **C.** Lure discrimination index (LDI) indicating bias towards pattern separation (positive) or completion (negative). Group distributions as un-mirrored violin plots (probability density functions), boxplots with 1st, 2nd (median), and 3rd quartiles, whiskers with 2nd and 98th percentiles, and individual (vertically jittered) data points (Allen et al., 2019). Zero denotes the chance level of the respective measure; above-chance performance (larger proportion of correct responses than incorrect responses in the respective measure) is indicated by positive values and below-chance performance by negative values.

One aspect of IM is lure discrimination (LDI; e.g., Ngo et al., 2018; Toner et al., 2009; Figure 4C). A high LDI is achieved by a large proportion of trials in which lures could be identified as similar and a low proportion of trials in which lures were mistaken as old, indicating pattern separation. A lower score, in contrast, indicates greater generalization or a bias towards pattern completion (cf. Keresztes et al., 2017). A one-way ANOVA contrasting all groups revealed significant age differences (*F*(2,112) = 19.86, *p* < 0.001). Post-hoc *t*-tests showed no significant differences between children and young adults (*t*(58) = 0.82, *p* = 0.414) whereas both children (*t*(74) = 4.76,*p* < 0.001) and young adults (*t*(92) = 5.49,*p* < 0.001) showed a higher LDI than older adults did. Thus, whereas all groups generally exhibited a bias towards pattern separation (i.e., positive values), older adults were driven more towards pattern completion than children and young adults.

To control for the different task versions (see Experimental design), 23 additional young adults performed the shorter children’s task. In the shorter task, children and young adults did not differ in either IM (*t*(42) = –1.12, *p* = 0.267), CMO (*t*(42) = –1.15, *p* = 0.257), or LDI (*t*(42) = –1.58, *p* = 0.121), whereas young adults with the shorter task performed better than young adults with the longer task (IM: *t*(60) = 3.94,*p* < 0.001; CMO: *t*(60) = 2.96,*p* = 0.004; LDI: *t*(60) = 3.14, p = 0.003), indicating that children’s high performance was largely due to lower task difficulty.

### 3.2 Repetition effects in ERPs

Using paired sample *t*-tests (first level) and non-parametric cluster-based permutation analysis (second level), we derived group-specific time–electrode clusters that indicate the trial time points and electrodes in which first and second object presentations showed reliable differences. We considered clusters whose test statistic exceeded the 97.5th percentile for its respective reference distribution as showing reliable repetition effects.

With this procedure, we identified two clusters in children, three in younger adults, and four in older adults (all clusters *p* ≤ 0.002) with broadly overlapping topography and latency (see Figure 5 for details). For all age groups, there was a cluster over posterior electrodes showing lower positive amplitudes for the second versus first stimulus presentation (i.e., repetition suppression; RS1; Figure 5A). This cluster appeared earliest in children (380–868 ms after stimulus onset), slightly later and with shorter duration in young adults (492–840 ms), and latest and with the shortest duration in older adults (568–884 ms). Within similar time windows another repetition suppression cluster (RS2) was identified at frontal and central electrodes which showed reduced negativity for repeated stimuli (Figure 5B). This cluster also appeared earliest in children (192–864 ms), followed by young adults (224–756 ms), and latest, as well as with much shorter duration in older adults (676–848 ms). For all age groups, both of these suppression effects occurred mainly while the ERP deflections came back to baseline, starting right at or after the peak in children and starting only shortly before returning to baseline in older adults. As the effects appeared at similar times but at different locations, and with opposite polarity, they could be two reflections of the same repetition suppression effect (perhaps resulting from the two sides of the dipole). To test this, we assessed whether the magnitudes of the effects correlated across subjects (see below).

**Figure 5.**
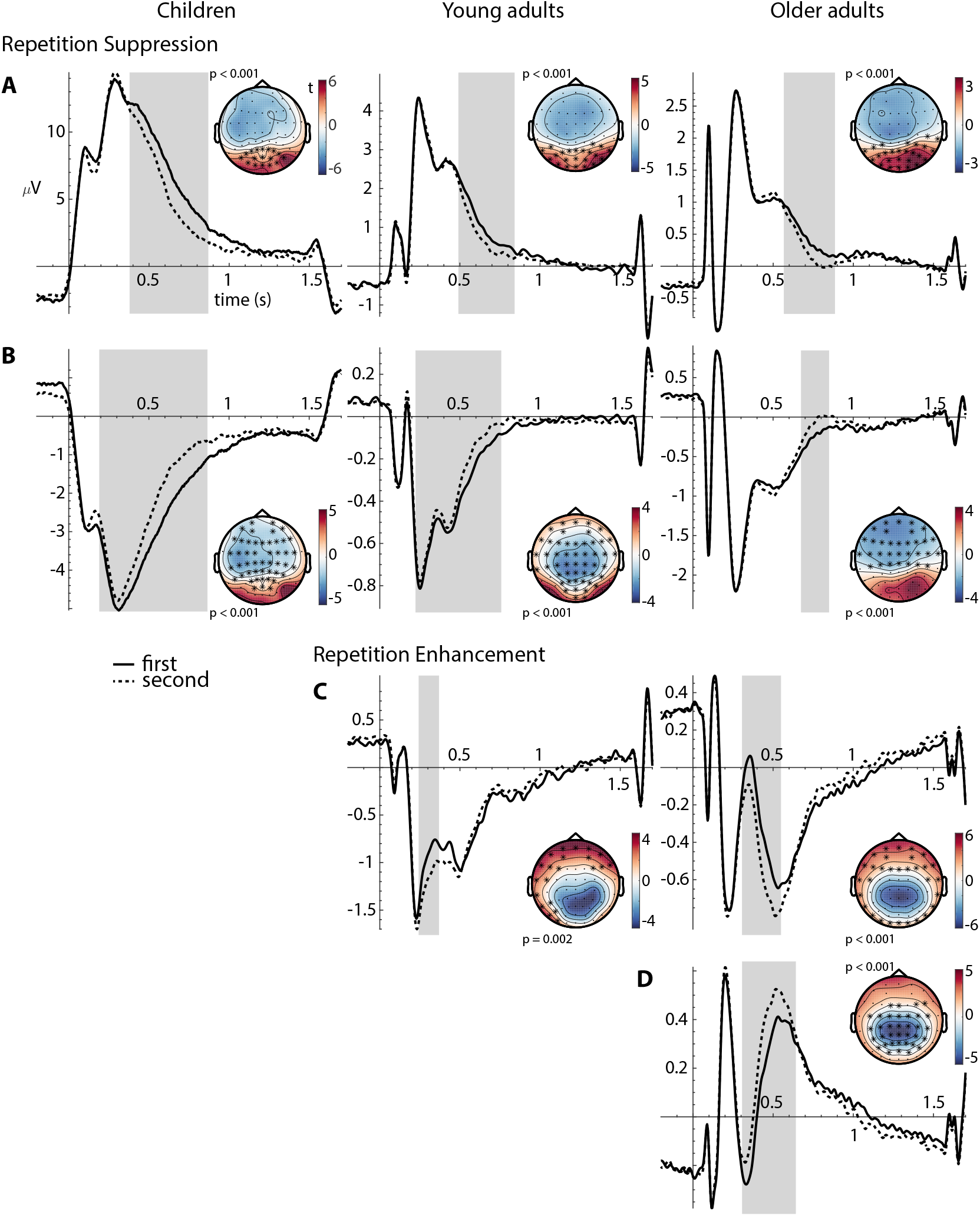
Group-specific repetition suppression (A, B) and repetition enhancement (C, D) effects. A. Positive posterior suppression effect (RS1). B. Negative fronto-central suppression effect (RS2). C. Negative frontal enhancement effect (RE1). D. Positive centro-parietal enhancement effect (RE2). ERPs are averaged over all trials in which objects were shown for the first time (solid line) or for the second time (dashed line) for children (left), young adults (middle), and older adults (right). The *x*-axis shows trial time (s) with stimulus onset at 0 (origin) and offset at 1.5 s, the *y*-axis shows amplitude *(μV)* with negative values plotted downwards. The time windows in which reliable differences between first and second presentation were identified (cluster-based permutation analysis) are shaded in gray. All ERPs are averaged over the respective electrodes in which the effects were identified, highlighted by asterisks in the respective topographical distributions plotted next to the ERPs. Topographies show the resulting *t*-values from contrasting ERPs of first and second presentations, averaged over the respective significant time windows. The *p*-values from the clusterbased permutation analysis are provided for each time–electrode cluster.

Furthermore, we identified a repetition enhancement effect (RE1) for both young and older adults that showed stronger (more negative) activity for the second than for the first object presentations over mainly frontal and temporal electrode sites (young adults: 244–368 ms; older adults: 308–548 ms; Figure 5C). An opposite enhancement effect (RE2) was only identified for older adults over centro-parietal regions at 308– 640 ms after stimulus onset (Figure 5D). In analogy to the two suppression effects, the opposite enhancement clusters identified in older adults could be two reflections of the same effect, which would be indicated by a high correlation between the effect sizes.

To test for correlations between the repetition effects, we extracted the individual */*-values (first-level analysis) from the identified time–electrode clusters (second-level analysis) and averaged them across the respective time points and channels for each subject. These *t*-values were then correlated across subjects. High correlations would indicate that participants who showed large effect sizes in, for example, RS1, also showed a large RS2 effect, indicating that they are not two independent effects potentially serving different functions. Indeed, for all age groups as well as across groups, the magnitudes of the RS1 and RS2 effects correlated strongly (children: *r* = 0.86, *p* < 0.001; young adults: *r* = 0.72, *p* < 0.001; older adults: *r* = 0.81, *p* < 0.001; across groups: *r* = 0.81, *p* < 0 .001; Pearson correlation). Likewise, the two enhancement effects of the older adults were highly correlated (*r* = 0.72, *p* < 0.001). Hence, we pooled the repetition effects into composite RS and RE effects by averaging the respective effect sizes for each subject. RS and RE effects did not correlate, potentially indicating two functionally different effects.

### 3.3 Age differences in ERP repetition effects

Repetition effects were identified for all age groups (see Figure 5). In a next step, we examined whether the size of the effects differed between age groups (see Figure 6 left). Because of overall age differences in EEG amplitudes, rather than computing and comparing the raw amplitude difference between the ERPs to first and second presentations, it is more appropriate to compare the effect sizes, i.e., the respective *t*-values of the contrast, which are independent of the overall amplitude. For group comparisons, *t*-values were transformed into *z*-values.

**Figure 6.**
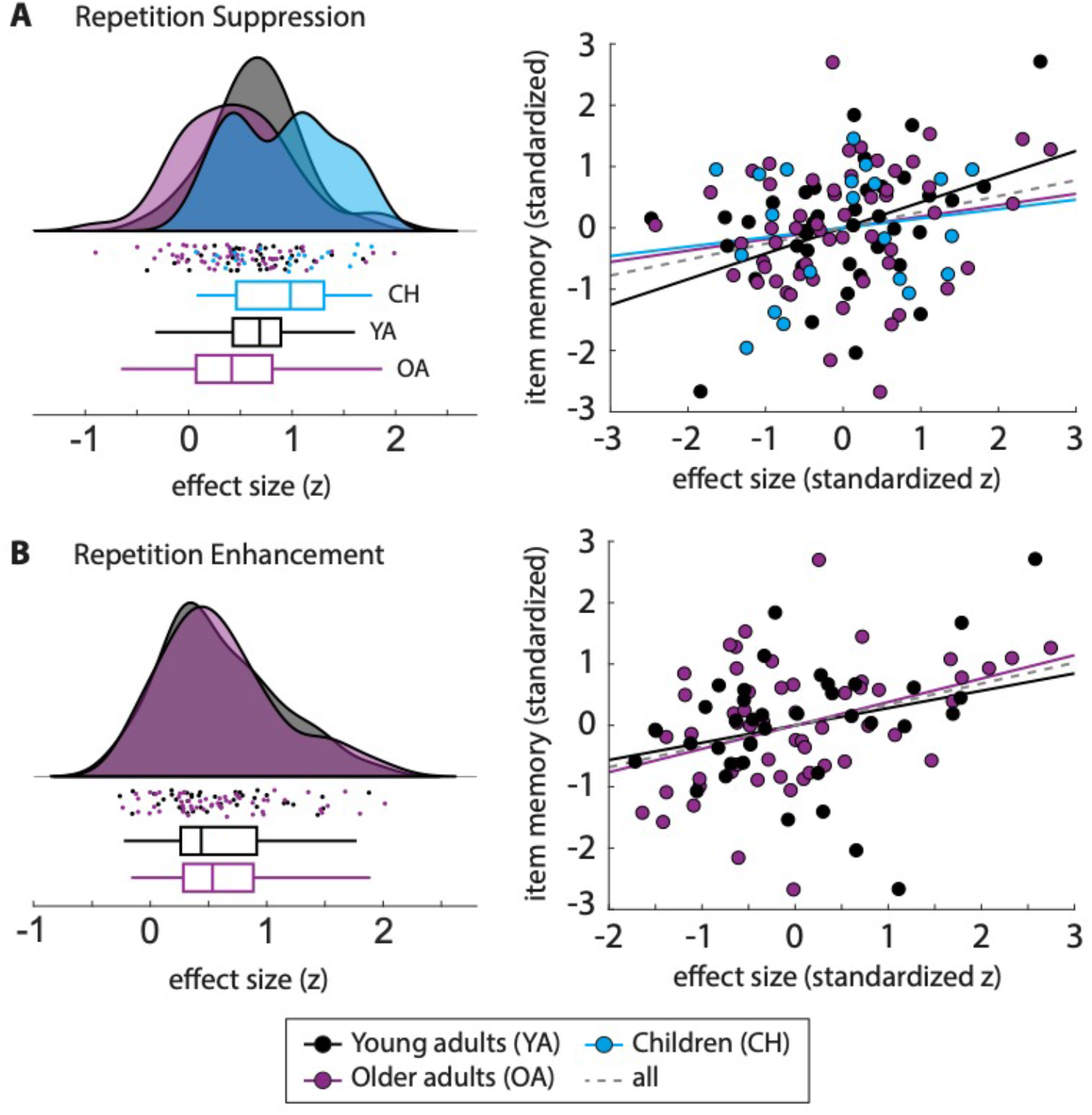
Repetition suppression (A) and enhancement (B) effect sizes for each group (left) and correlations between effect sizes and item memory performance (right). Left: Group distributions as un-mirrored violin plots (probability density functions), boxplots with 1st, 2nd (median), and 3rd quartiles, whiskers with 2nd and 98th percentiles, and individual (vertically jittered) data points (Allen et al., 2019) of the mean effect sizes (*z*-transformed *t*-values within the respective clusters) in children (blue), young adults (black), and older adults (purple). Right: Association (Pearson correlation) between individual standardized effect sizes (*z*-transformed *t*-values; *x*-axis) and standardized item memory performance (IM; *y*-axis) indicated by least-squares lines, separately for each group (colors as above) and across groups (gray dashed line). For statistics and correlations with other performance measures, see Table 2.

The composite RS effect sizes (*z*-values) differed between the age groups (*F*(2,112) = 5.72, *p* = 0.004; one-way ANOVA) with children showing larger effects than adults (children vs. young adults: *t*(58) = 2.32, *p* = 0.024; children vs. older adults: *t*(74) = 3.11, *p* = 0.003; post-hoc *t*-tests) and no differences between young and older adults (*t*(92) = 1.47, *p* = 0.145) (see Figure 6A left). The enhancement effects identified for young adults (RE1) and older adults (composite of RE1 and RE2) did not differ in their effect sizes (*t*(92) = −0.11, *p* = 0.915) (see Figure 6B left).

### 3.4 Repetition effects are positively linked to memory performance

We examined the relationship between the individual mean effect sizes of the composite repetition effects (see above) and the individual performance measures (see Behavioral analysis) by modelling their correlations using maximum likelihood estimates, (a) separately for each age group, and (b) across all groups. Because there were overall age differences in both ERP effects and memory performance (see above), we standardized both measures within groups to test for the brain–behavior association independent of age differences. The resulting estimated correlation coefficients are presented in Table 2.

**Table 2:** Estimated correlation coefficients (*r*) for modelled correlations of individual ERP effect sizes (standardized *z*-transformed *t*-values) of the composite repetition suppression (RS) and repetition enhancement (RE) effects with memory performance measures (IM, CMO, LDI; standardized) for all age groups separately as well as across groups (All). Correlations for item memory (IM) are also plotted in Figure 6. Likelihood ratio (LR) tests were used to compare the models of group-specific versus group-independent brain–behavior correlations. Significant LR would indicate that correlations differed between age groups.

| Memory | Effect | Children | Young adults | Older adults | All | Model comparison |  |  |
| --- | --- | --- | --- | --- | --- | --- | --- | --- |
| | | $r$ | $r$ | $r$ | $r$ | $LR$ | $df$ | $p$ |
| IM | RS | 0.15 | 0.42 | 0.19 | 0.26 | 1.98 | 2 | 0.372 |
|  | RE |  | 0.28 | 0.38 | 0.34 | 0.32 | 1 | 0.572 |
| CMO | RS | -0.26 | 0.07 | 0.00 | -0.02 | 1.56 | 2 | 0.459 |
|  | RE |  | -0.03 | 0.29 | 0.16 | 2.58 | 1 | 0.108 |
| LDI | RS | -0.01 | 0.35 | -0.07 | 0.08 | 4.44 | 2 | 0.109 |
|  | RE |  | 0.32 | -0.02 | 0.12 | 2.85 | 1 | 0.091 |

*IM = item memory, CMO = category memory only, LDI = lure discrimination index, RS = repetition suppression, RE = repetition enhancement, LR = likelihood ratio, df = degrees of freedom*

Across groups, the composite RS effect correlated significantly with IM (see Figure 6A right) in the direction that larger RS were associated with better memory performance. The RE effect that only occurred in adults also showed a positive association to IM across young and older adults (Figure 6B right). In contrast, CMO and LDI did not show significant brain–behavior correlations across groups (see Table 2).

We tested whether the models (a and b) differed in their model fits using likelihood ratio (LR) tests. For all repetition effects (RS, RE) and performance measures (IM, CMO, LDI), the model fits were not significantly different from each other (see Table 2). That is, the model that allowed age group-specific brain–behavior relations did not explain the data significantly better than the model with fixed correlations across groups. This indicates that the brain–behavior associations did not differ across the age groups. Therefore, in the following we do not interpret the group-specific but only the cross-group brain–behavior correlations.

## 4. Discussion

The present study investigated whether processes of memory formation, as reflected in neural repetition effects, are associated with inter-individual differences in item and category memory. Specifically, we were interested whether age group differences in memory performance would show as age-differential associations between repetition effects and memory specificity in a lifespan sample. For this, we compared repetition-related differences in ERPs during incidental object encoding in groups of 7–9-year-old children, 18–30-year-old young adults, and 65–76-year-old older adults in relation to their later recognition performance. We distinguished between memory for specific items and for the general stimulus category only. We identified reliable ERP repetition suppression effects for all age groups and repetition enhancement among adults. Across groups, item-specific memory performance was positively associated with the magnitude of repetition suppression and enhancement. In sum, we demonstrate common neural mechanisms of memory encoding across the lifespan.

Overall, we found little evidence for age differences in memory performance. While children showed better item-specific memory compared with adults who performed a longer task, there were no performance differences between children and a control sample of young adults who also performed the children’s task version. With regard to simple recognition, this is in accordance with previous findings also showing no age differences in object recognition between children (>6 years old), adolescents, and young adults (Golarai et al., 2007; Rollins and Cloude, 2018). With regard to memory specificity as assessed by versions of the mnemonic similarity task (MST; Stark et al., 2013, 2019), Rollins and Cloude (2018) observed lower lure discrimination performance in children aged 5-9 (but not aged 11-12) than young adults. However, using a similar task, Ngo et al. (2018) only found poorer lure discrimination in 4-year-olds but not in 6-year-olds compared to young adults, which is in line with the current findings of adult-like lure discrimination in 7-9-year-olds. Overall, although brain maturation has shown to proceed well beyond childhood, especially regarding hippocampal (Keresztes et al., 2018, 2017) and prefrontal regions (Bunge et al., 2002), our current findings substantiate the evidence that in specific task settings such the ones used in the current study, older children (here, 7–9-year-olds) may be already able to reach adult-like item-specific memory and lure discrimination performance.

At the other end of the lifespan, also younger and older adults did not differ with respect to recognition memory, which is in accordance with previous work showing much less pronounced or no age differences for item compared to associative memory, especially under incidental encoding conditions (Old and Naveh-Benjamin, 2008). However, one might have expected that older adults would tend to generalize more and thus show worse item-specific memory than young adults, perhaps accompanied by greater reliance on category memory (Koutstaal and Schacter, 1997; Tun et al., 1998). Although this was not the case in these aggregated performance assessments, evidence in this direction is nevertheless provided by age-related differences in lure discrimination. The ability to recognize lures as “similar” and not “old” is only achievable if the original targets have been encoded with high specificity. Here, young adults outperformed older adults, which corroborates previous findings (Schacter et al., 1997; Stark et al., 2015, 2013; Stark and Stark, 2017) and has been shown to be related to age-associated disruptions of the intra-hippocampal circuitry (Shing et al., 2011; Wilson et al., 2006; Yassa et al., 2011).

All in all, the behavioral age pattern shows that 7–9-year-old children may under specific circumstances already show adult-like levels of item-specific recognition and lure discrimination. Further, while aggregated performance measures did not show differences between young and older adults, some measures (e.g., lure discrimination) did, indicating aging-related behavioral disparity in specific memory aspects.

With regard to our ERP results, we showed that all age groups exhibited repetition-related changes in the neural responses during encoding, suggesting common neural mechanisms of memory formation overall. We identified repetition suppression (for all age groups) and repetition enhancement (only for adults) with overlapping topographies and time windows across age groups but age-related differences with regard to the magnitudes of the effects.

The topography and timing of the observed repetition effects are in line with the effects described in the ERP literature that have been observed in adult samples (cf. Henson et al., 2004; Lawson et al., 2007; Penney et al., 2001; Rugg and Doyle, 1994; Stefanics et al., 2018). The repetition suppression effects may be linked to modulations in the P600 ERP component, which is sensitive to (word) frequency and familiarity (Rugg and Doyle, 1994; Rugg 1990), and in FN400, which is also associated with familiarity (in adults; Curran, 2004; Friedman and Johnson, 2000; Mecklinger, 2000). Thus, modulations in these components may suggest differences in stimulus familiarity that are certainly to be expected when items are repeated. In children, familiarity-sensitive FN400 has also been observed before albeit less consistently (Boucher et al., 2016; Congdon et al., 2012; Haese and Czernochowski, 2016; Mecklinger et al., 2010). For example, Boucher et al. (2016) found old/new repetition effects in the FN400 component of school children for both concrete and abstract object images. Since we observed these modulations in all age groups, item familiarity during the second item presentation may be related to suppressed neural activity for children, young, and older adults alike, which is in line with their comparable item recognition abilities.

The repetition enhancement effect that we observed only for adults partially resembles the original ERP repetition effect described in early studies (Rugg and Doyle, 1994). It has been argued that this reflects modulations in several ERP components, mainly a decrease in N400, which is related to priming and contextual integration, and an increase in P300, which is related to working memory and stimulus categorization (Friedman, 2000; Lawson et al., 2007; Rugg and Doyle, 1994; Van Petten and Senkfor, 1996). The observation that the ERPs of repeated stimuli show changes in these components may indicate successful integration of the items into internal object representations. The finding that children did not show such an effect, while young and older adults did not differ in their enhancement effects, may suggest that the neural mechanisms of forming representations is not yet fully mature in children, but still intact in older adults. This would be largely consistent with previous studies showing developmental differences in context integration and binding in general (Káldy and Kovács, 2003; Raj and Bell, 2010) and also in relation to working memory (Fandakova et al., 2014). An alternative interpretation of repetition enhancement and suppression effects has been put forward by Henson et al. (2000): They suggested that repetition enhancement may reflect the formation of new representations of previously unfamiliar stimuli, whereas suppression may reflect the strengthening of already existing representations (see also Grossmann et al., 2009). According to this interpretation, one would assume that children have had less experience with the presented objects (cf. Brod et al., 2013) and therefore should show not less, but more enhancement than adults, who presumably had previously well-established representations of all the objects. Our data does not support such an interpretation. Nevertheless, the absence of repetition enhancement in children may also be due to deficient statistical power as the children’s group was smaller and they had fewer trials than the adult groups. Since other studies on repetition effects in children of this age are lacking, we can only speculate on whether children may indeed exhibit no enhancement. If this is the case, it may indicate that the underlying neural mechanisms related to context integration and working memory are not yet fully developed, which, however, did not appear to have impaired children’s memory performance.

While similar patterns of suppression effects occurred in all three age groups, we observed differences in the magnitudes of the effects between groups. Comparing the composite suppression effect between age groups yielded the largest effect sizes for the children’s group. Young and older adults differed in neither the magnitudes of suppression nor enhancement, which is generally in line with previous studies (e.g., Hamberger and Friedman, 1992; but see Friedman et al., 1993). On this between-group level, the cooccurrence of children’s strong memory performance and larger repetition effects while the adult groups differed in neither, may suggest that memory success is associated to strong repetition suppression, in line with the hypothesis that repetition effects reflect successful memory formation. Accordingly, when we examined the relation of individual differences in repetition effects and memory on the item- and category-level both within and across age group, we found a positive relationship of the extent to which neural activity is altered in response to repeated input and individual explicit memory performance. Across groups, individuals exhibiting larger repetition effects during encoding showed better item recognition memory than individuals with smaller repetition effects did. Importantly, this was not due to overall age group differences, such as children showing both larger effects and better performance (see above), but the association remained when overall age differences were eliminated by standardizing the measures within groups. Furthermore, the estimated brain–behavior associations did not significantly differ between the groups, suggesting largely common neural correlates of successful memory encoding across the lifespan. The finding that the repetition effects were linked to specific item memory and not to mere category memory furthermore suggests that repetition-sensitive neural responses reflect highly detailed memory encoding. Moreover, repetition effects were not related to mere lure discrimination but specific item memory, including correct target recognition, lure detection, and foil rejection, indicating that the LDI alone may provide an incomplete picture with regard to the abilities reflecting highly specific memory representations. All in all, we show that ERP repetition effects provide an age-independent indicator of individual differences in successful encoding of item-specific details.

To conclude, we found that memory formation processes that manifest in repetition-sensitive ERP components are related to individual differences in item recognition performance across the lifespan. The effects did not differ between age groups, indicating age-independent reflections of memory encoding that have not been shown before. These novel findings highlight the significance of encoding mechanisms that facilitate the formation of high-quality memory representations and enable highly specific recognition and item distinction.

## Acknowledgements

This research was conducted within the projects Lifespan Age Differences in Memory Representations (LIME) (PI: MCS) and Lifespan Rhythms of Memory and Cognition (RHYME) (PI: MWB) at the Max Planck Institute for Human Development, Germany, and the Department of Vision, Visual Impairments & Blindness (PI: SW) at TU Dortmund University’s Faculty of Rehabilitation Sciences, Germany. LM and SW thank the Research Department of Neuroscience at Ruhr-University Bochum for access to EEG equipment and analysis software. VRS was a fellow of the International Max Planck Research School on the Life Course. SW was supported by a grant from the Volkswagen Foundation (Lichtenberg Professorship, 97 097). MWB was supported by grants from the German Research Foundation (DFG; WE 4269/2-1 and WE 4269/5-1) and an Early Career Research Fellowship by the Jacobs Foundation. MCS was supported by the MINERVA program of the Max Planck Society. We thank Gabriele Faust, Aleksandr Merkulov, and all student assistants who helped with organization and data collection, Katharina Limbach for her support in data preprocessing, Julia Delius for her editorial assistance, all members of the projects for their helpful feedback, and all study participants for their time.

## Supplemental material

### Encoding conditions

The 80 object categories were equally divided into four conditions (see Figure S1) that differed with respect to (a) the number of different presented exemplars from one category (either two or four), and (b) the number of exemplar repetitions (either two or four times). This resulted in four encoding conditions: (1) Baseline (BL) condition with two exemplars per category that were each presented twice, (2) High-Repetition (HR) condition with two exemplars per category that were each presented four times, (3) High-Exemplar (HE) condition with four exemplars per category that were each presented twice, (4) High-Repetition-and-Exemplar (HRE) condition with four exemplars per category that were each presented four times.

Twenty object categories were randomly selected for each condition. For those conditions with two exemplars per category (BL and HR), two of the four available exemplars were randomly selected for presentation. Since the number of exemplars and repetitions differed between conditions, the number of trials in each condition varied accordingly: 80 in the BL condition, 160 in the HR condition, 160 in the HE condition, and 320 trials in the HRE condition. This adds up to a total of 720 trials. A shorter version of the task was used for the children. They performed only the first half of the adult task, comprising 40 distinct object categories, i.e., 10 in each encoding condition, which results in 360 experimental trials (40 BL, 80 HR, 80 HE, 160 HRE).

For those categories that belonged to an encoding condition with four exemplars (HE and HRE), two of these were randomly selected to be tested in the recognition task (as old items).

Since we did not have specific hypotheses how these encoding conditions would affect event-related repetition effects, for the purpose of the analysis presented in this article, we collapsed across all conditions and focused on the comparison of the first and second presentation of an object, independent of its condition.

**Figure S1.**
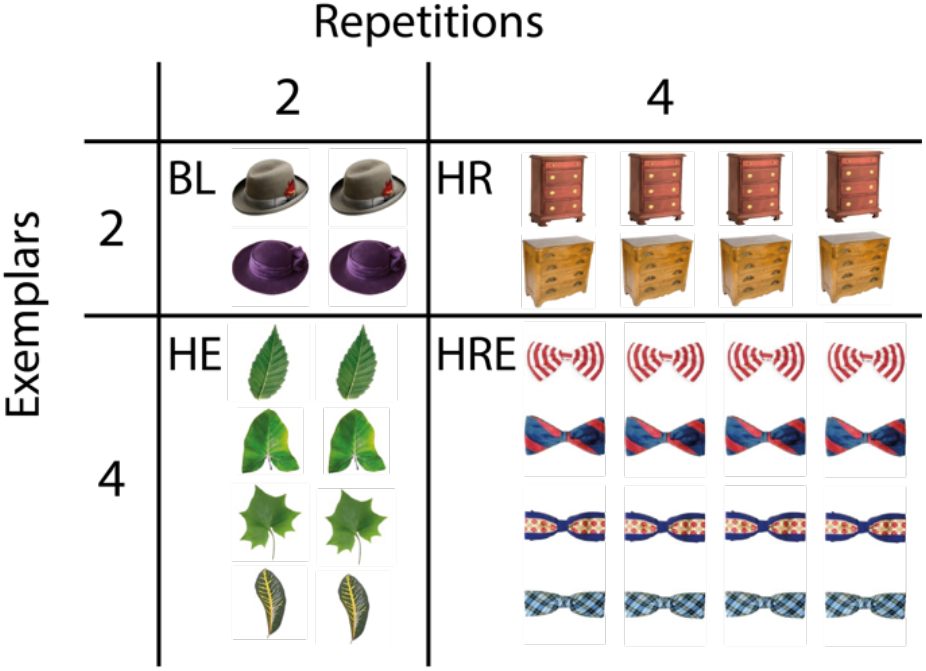
Encoding conditions. The number of item repetitions (2 or 4) and category exemplars (2 or 4) were manipulated during encoding, resulting in 4 encoding conditions: Baseline (BL), High-Repetition (HR), High-Exemplar (HE), and High-Repetition-and-Exemplar (HRE). Each condition comprised 20 object categories for adults, and 10 object categories for children. One sample category per condition is shown for illustration.

### The effect of repetitions on item memory

In the following, we asked whether item and category repetitions might affect memory performance differentially in children, younger adults, and older adults. We initially expected better item-specific memory performance when items were repeated more often and worse performance when more exemplars were presented during encoding. This was based on the hypothesis that more repetitions would provide more opportunity to encode item-specific details leading to both better target recognition as well as rejection of similar lures (Benjamin, 2001; but see Reagh and Yassa, 2014), whereas more exemplars would create stronger interference, making it harder to distinguish between similar items (Anderson, 2015). The latter is supported by previous findings showing that increasing the number of exemplars from the same object category, reduced participants’ ability to discriminate between lures and targets from that category (Gallo, 2004; Konkle et al., 2010; Omohundro, 1981; Poch et al., 2019). Alternatively, interference from similar memories can also trigger a repulsion of memory representations, suggested by the findings of less overlapping activation patterns in the hippocampus (pattern separation), which was associated with less memory interference, i.e., better performance (Chanales et al., 2020, 2017; Favila et al., 2016). That is, both scenarios – better or worse item memory when more exemplars were presented – are conceivable.

To examine these predictions, we conducted a three-way mixed ANOVA with age group as the between-subjects factor, number of repetitions and number of exemplars as within-subjects factors, and item memory performance as the dependent variable. The results revealed a main effect of age (*F*(2,109) = 7.48, *p* < 0.001) with the same pattern of age differences as reported above, and a main effect of the number of exemplars (*F*(1,109) = 8.53, *p* = 0.004). Post-hoc t-tests demonstrated better item memory when four rather than two exemplars were presented (*t*(111) = 3.58,*p* < 0.001), revealing a beneficial effect of a higher number of category exemplars. This is in line with the repulsion prediction (see above), whereas the number of repetitions did not influence item memory (*F*(1,109) = 0.99, *p* = 0.321). To better disentangle the effect of repetitions and exemplars for item-specific memory, we further examined the potential effect of repetitions by directly contrasting memory performance for items from the HR and BL condition, i.e., independently of the exemplar manipulation. Across age groups, this revealed better item memory for the HR condition (*t*(111) = 2.72, *p* = 0.008; paired-sample *t*-test).

Since there were no interactions between age and item manipulations, we largely disregard the encoding conditions and do not separate the trials accordingly in the analyses presented here.

### Lifespan differences in ERPs

The ERPs of children, young adults, and older adults at selected representative electrode sites are shown in Figure S1. Children exhibited overall higher amplitudes than adults, which is consistent with previous findings in the literature (Coch et al., 2005; Dustman and Beck, 1969; Mueller et al., 2008). This could, for example, be due to differences in skull thickness (Frodl et al., 2001) but in the current study it may also be a result of the different EEG systems and laboratories used to test the children and adults.

**Figure S2.**
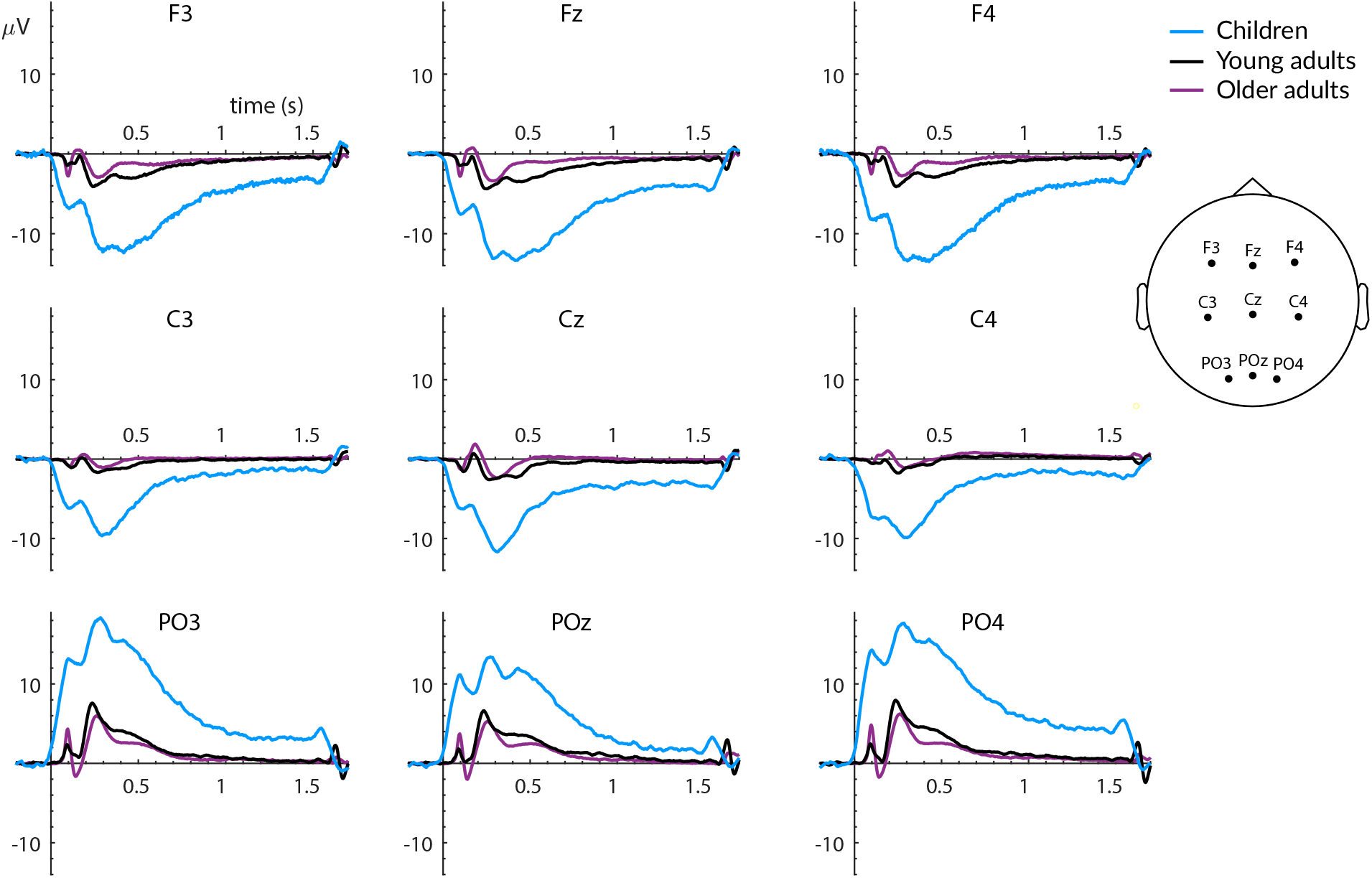
Event-related potentials (ERPs) at selected electrode sites, averaged over all trials in which an object was shown for the first time, for all children (blue), young adults (black), and older adults (purple). The *x*-axis shows trial time (s) with stimulus onset at 0 (origin) and offset at 1.5 s, the *y*-axis shows amplitude *(μV)* with negative values plotted downwards. Data were baseline-corrected with an absolute pre-stimulus baseline of 200 ms.

### Repetition suppression for all presentations

**Figure S3.**
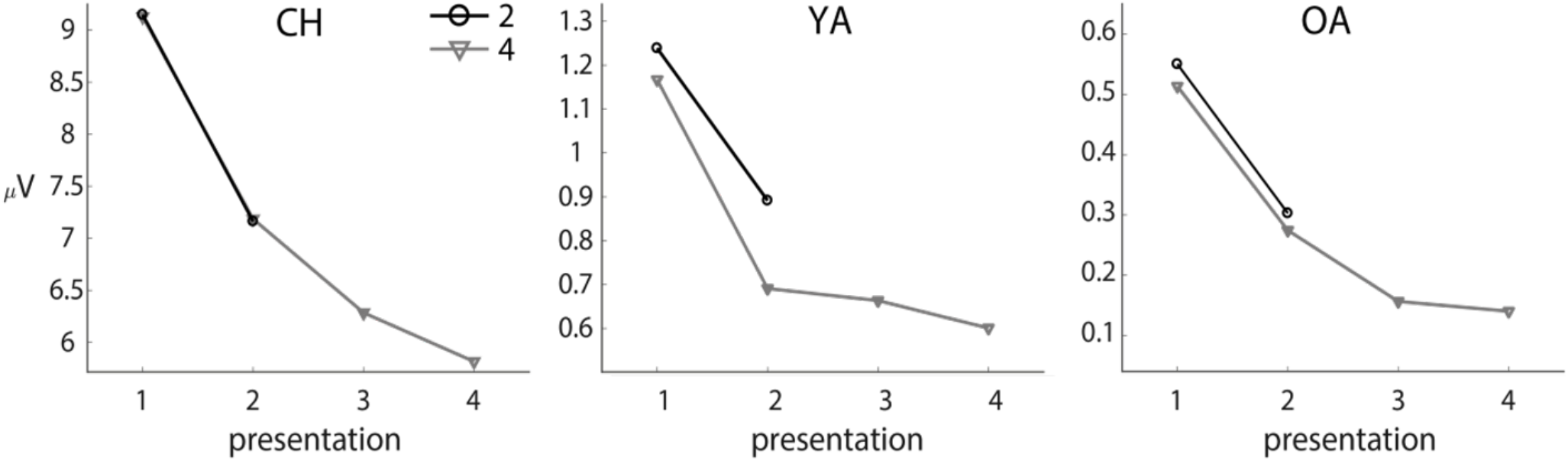
Mean amplitudes (μV; y-axis) within identified clusters (first repetition suppression effect) for all repeated presentations (x-axis), averaged over conditions with two repetitions (N and HE; black line) and conditions with four repetitions (HR and HRE, gray line) for children (CH; left), young adults (YA; center), and older adults (OA; right).

**Table S1.** Covariates

| Group | Processing speed<br>DSST<sup>1</sup> *M*(*SD*) | Verbal knowledge STW<sup>2</sup><br>*M*(*SD*) | Dementia screening<br>MMSE<sup>3</sup> *M*(*SD*) | Verbal digit span task<sup>4</sup><br>*M*(*SD*) |
| --- | --- | --- | --- | --- |
| Children | 29.62(7.00) | N/A | N/A | 13.43(2.04) |
| Young adults<br>(EEG) | 63.97(10.67) | 22.21(4.16) | N/A | N/A |
| Young adults<br>(control) | 70.2(6.3)* | N/A | N/A | 20.83(3.06) |
| Older adults | 47.58(8.99) | 28.72(3.30) | 28.95(1.21) | N/A |

*M* = mean, *SD* = standard deviation, <sup>1</sup> Digit Symbol Substitution Test (Wechsler, 1981): number of correct responses in 90 s, \* for the control young adults group, test time was mistakenly set to 120 s; the provided value is converted to responses / 90 s, <sup>2</sup> Spot-the-Word test (Mehrfachwahl-Wortschatz-Intelligenztest; Lehrl, 1977): number of correct responses, <sup>3</sup> Mini-Mental State Examination (Folstein et al., 1975); <sup>4</sup> Petermann and Petermann, 2011

